# VaxOptiML: Leveraging Machine Learning for Accurate Prediction of MHC-I & II Epitopes for Optimized Cancer Immunotherapy

**DOI:** 10.1101/2024.06.10.598389

**Authors:** Dhanushkumar T, Sunila B G, Sripad Rama Hebbar, Prasanna Kumar Selvam, Karthick Vasudevan

## Abstract

In the realm of cancer immunotherapy, the ability to accurately predict epitopes is crucial for advancing vaccine development. Here, we introduce VaxOptiML (available at https://vaxoptiml.streamlit.app/), an integrated pipeline designed to enhance epitope prediction and prioritization. Utilizing a curated dataset of experimentally validated epitopes and sophisticated machine learning techniques, VaxOptiML features three distinct models that predict epitopes from target sequences, pair them with personalized HLA types, and prioritize them based on immunogenicity scores. Our rigorous process of data cleaning, feature extraction, and model building has resulted in a tool that demonstrates exceptional accuracy, sensitivity, specificity, and F1-score, surpassing existing prediction methods. The robustness and efficacy of VaxOptiML are further illustrated through comprehensive visual representations, underscoring its potential to significantly expedite epitope discovery and vaccine design in cancer immunotherapy, Additionally, we have deployed the trained ML model using Streamlit for public usage, enhancing accessibility and usability for researchers and clinician.

## Introduction

Cancer, a complex and multifaceted disease, is characterized by the uncontrolled growth and proliferation of abnormal cells, driven by a combination of genetic, epigenetic, and environmental factors [1]. This aberrant cellular behavior disrupts the normal regulatory mechanisms governing cell division, differentiation, and apoptosis, leading to the emergence of various hallmarks such as sustained proliferative signaling, evasion of growth suppressors, and resistance to cell death [2].

At the molecular level, cancer is marked by genetic alterations that disturb the balance between oncogenes, promoting cell proliferation, and tumor suppressor genes, inhibiting tumor growth [3]. These alterations encompass mutations, chromosomal rearrangements, gene amplifications, and epigenetic modifications, collectively contributing to the dysregulation of key signaling pathways involved in cell cycle control, DNA repair, and apoptosis [3]. Moreover, cancer development and progression entail a dynamic interplay between tumor cells and the surrounding microenvironment, comprising diverse cell types, extracellular matrix components, and signaling molecules [5]. Tumor cells can manipulate the microenvironment to support their growth, evade immune surveillance, and facilitate metastasis [5]. Conversely, the microenvironment can exert both pro-tumorigenic and anti-tumorigenic effects, influencing cancer progression and treatment response [5].

Cancer is highly heterogeneous, encompassing a spectrum of tumor types with distinct molecular and clinical characteristics [6]. This heterogeneity poses challenges for diagnosis, prognosis, and treatment, as tumors may exhibit differential responses to therapy and varying metastatic potentials [6]. Advancements in molecular profiling technologies have enabled the elucidation of the molecular landscape of cancer, identifying key drivers of tumorigenesis and therapeutic targets [7]. The immune system plays a critical role in cancer surveillance and defense, with immune cells capable of recognizing and eliminating transformed cells through immunosurveillance [8]. However, tumors can evade immune detection and suppression by upregulating immune checkpoint molecules, inhibiting T cell activation and effector function [8]. Immunotherapy strategies aim to restore and enhance anti-tumor immune responses, offering promising outcomes in some cancer patients [9].

Nevertheless, challenges such as tumor heterogeneity, immune evasion mechanisms, and autoimmune toxicities hinder the success of immunotherapy [10]. Epitope-based vaccines represent a promising approach within cancer immunotherapy, stimulating anti-tumor immune responses by targeting specific antigenic peptides presented on major histocompatibility complex (MHC) molecules [11]. The selection of optimal MHC epitopes for cancer vaccines is crucial for eliciting robust and durable anti-tumor immune responses [12]. Machine learning (ML) has emerged as a powerful tool for epitope prediction, offering a data-driven approach to identify candidate epitopes with high specificity and immunogenicity [14].

In this study, we propose leveraging ML techniques to develop a novel model for predicting T cell epitopes in cancer vaccines, aiming to enhance the accuracy and robustness of epitope prediction for personalized immunotherapies [15].

## Methods and Materials

### Curated Cancer Epitope Dataset

We utilized a comprehensive database from the Cancer Epitope Database and Analysis Resource (CEDAR) [16], which stores experimentally validated cancer epitopes along with their source materials, ensuring the dataset’s reliability and accuracy. From this database, we extracted 3,803,742 epitope sequences, their starting and ending sites, UniProt IDs, and associated HLA alleles, focusing on epitopes of human origin, covering a wide range of cancer types. Additionally, we sourced 289,117 non-epitopes from the IEDB database [17].

### Data Cleaning and Feature Extraction

To ensure data quality, we removed duplicate values and considered only human HLA alleles, filtering out 200 unidentified IDs. Using the UniProt and NCBI IDs associated with each epitope, we retrieved full-length protein sequences, providing comprehensive information about the parent proteins [18]. Using Biopython libraries, specifically Bio.SeqUtils.ProtParam, we extracted various physicochemical properties of the epitopes and protein sequences. These properties offered insights into their biochemical characteristics [19]. We computed the percentage composition of each amino acid in both epitopes and protein sequences, influencing antigenicity and binding affinity. Additionally, we calculated the total number of atoms for oxygen, carbon, hydrogen, nitrogen, sulfur, and the overall atom count in both epitopes and protein sequences. These atomic composition features helped understand the chemical and structural properties of the sequences ([20] A visual representation of the dataset with their features used in this analysis is shown in Figure 2. In total, we derived 46 distinct features for the epitopes and 45 features for the corresponding protein sequences, culminating in a comprehensive set of 91 features, as detailed in Supplementary Table 1. Epitopes were labeled as Target 1 and non-epitopes as Target 0. We addressed null values through imputation methods tailored to the data’s skewness, ensuring minimal bias and maximum data integrity. Rigorous outlier detection techniques using pandas identified and removed 150 outliers, enhancing the overall quality and reliability of our dataset [21, 22]

**Table 1.** Results of accuracy and other performance metrics for ensemble classifier model.

| Models | Accuracy | Sensitivity | Specificity | Precision | F1 score | AUC |
| --- | --- | --- | --- | --- | --- | --- |
| Bagging Classifier, | 0.765 | 0.757 | 0.770 | 0.770 | 0.798 | 0.755 |
| Extra Trees Classifier | 0.761 | 0.850 | 0.646 | 0.756 | 0.800 | 0.748 |
| Random Forest Classifier | 0.760 | 0.834 | 0.667 | 0.763 | 0.797 | 0.750 |

**Figure 1.**
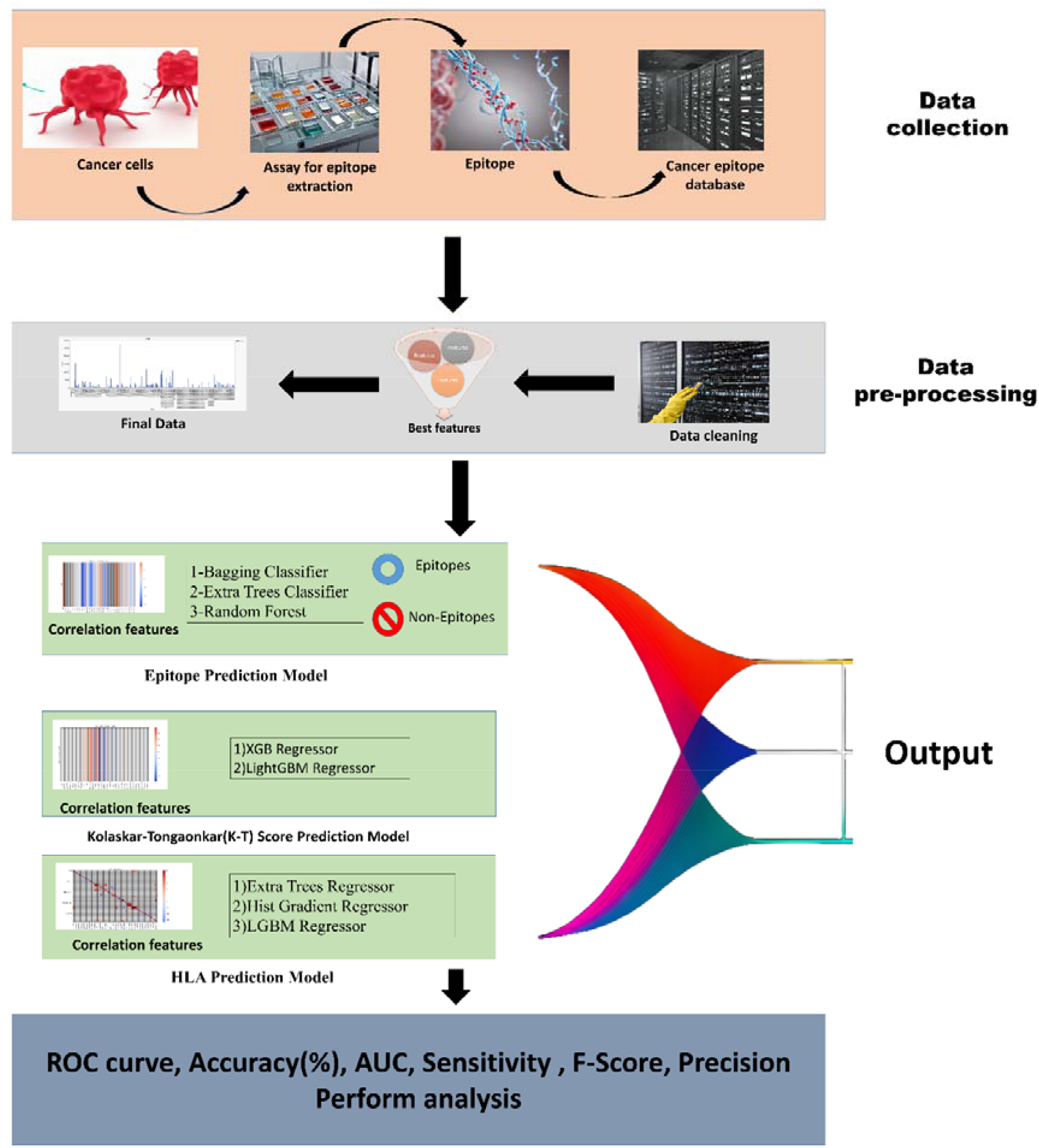
Schematic representation of methodology used to build the model

**Figure 2.**
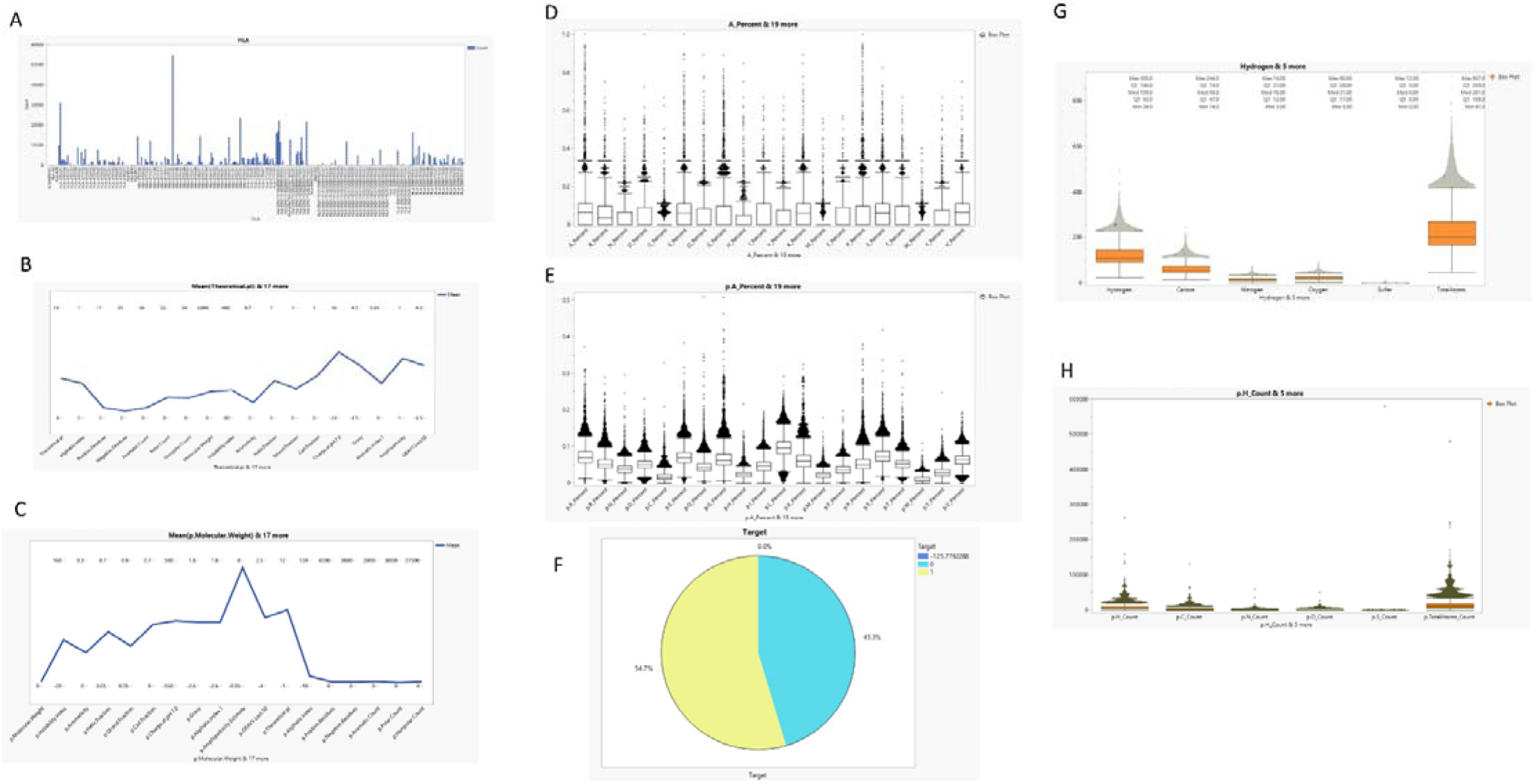
A) Bar graph of total HLA’s used in this dataset, B) Physio-chemical properties of epitopes, C) Physio-chemical properties of Protein sequence, D) Amino acid composition of epitopes, E)Amino acid composition of Protein sequence, F) Percentage of Target Column in the dataset, G) Atomic composition of epitopes, H) Atomic composition of Protein sequence

### Kolaskar-Tongaonkar Score

We extracted the Kolaskar-Tongaonkar antigenicity score for all epitopes [21]. This score predicts the antigenic determinants (epitopes) on proteins, essential for designing vaccines and understanding immune responses. The method utilizes physicochemical properties of amino acid residues and their frequencies of occurrence in known epitopes. It works on the principle that certain residues tend to occur more frequently in epitopes than in non-epitope

## Model Building and Evaluation

### Epitope Prediction Model

For epitope prediction, we conducted a correlation analysis to identify the most correlated features in the dataset. We utilized three classifiers: the Bagging Classifier, the Extra Trees Classifier, and the Random Forest Classifier [23-25], chosen for their effectiveness in handling classification tasks and complex datasets. The dataset was prepared by selecting numerical features and excluding the target variable. Each classifier was evaluated based on their performance in predicting epitopes.

### HLA Prediction Model

To predict HLA epitopes, we conducted a correlation analysis to identify pertinent features within the dataset. We employed three algorithms: the Extra Trees Regressor, the Hist Gradient Regressor, and the LGBM Regressor [26-28]. We transformed HLA allele strings into numerical scores via a bespoke Python function, calculate_numerical_score, mapping amino acid sequences from HLA alleles to their respective hydrophobicity values.

### Kolaskar-Tongaonkar (K-T) Score Prediction Model

We utilized backward feature selection to predict K-T scores for epitopes, retaining influential features while eliminating less informative ones. A correlation analysis identified relevant features, guiding the selection process. For model implementation, we opted for the XGB Regressor and LightGBM Regressor [29-30], chosen for their effectiveness in capturing complex data patterns.

## Results

### Model Building and Evaluation

#### Epitope Prediction Model

For epitope prediction, we conducted a correlation analysis to identify the most correlated features in the dataset, which were then listed in Supplementary Table 2 (Figure 3). We utilized three classifiers: the Bagging Classifier, the Extra Trees Classifier, and the Random Forest Classifier[23-25], chosen for their effectiveness in handling classification tasks and complex datasets. The Bagging Classifier is known for reducing variance and avoiding overfitting by averaging predictions from multiple models trained on random subsets of the data. The Extra Trees Classifier enhances computational efficiency and manages high-dimensional data by constructing trees with randomly split nodes. The Random Forest Classifier, known for its robustness and accuracy, reduces overfitting by averaging multiple decision trees, improving generalization and providing insights into feature importance. The dataset was prepared by selecting numerical features and excluding the target variable. Each classifier was evaluated based on their performance in predicting epitopes, with the Bagging Classifier achieving the highest accuracy of 76.51%, Accuracy, Sensitivity, Specificity, Precision F1, score, and AUC scores of the models are tabulates in table 1 . The effectiveness of these classifiers in handling complex datasets and providing reliable predictions underscores their selection for this analysis. Bar graphs of the models with their ROC plots are shown in Figure 4, demonstrating their comparative performance.

**Table 2.** Results of accuracy and other performance metrics for ensemble regressor model’s (for HLA prediction).

| Metric | Extra tree regressor | Hist gradient regressor | Lgbm regressor |
| --- | --- | --- | --- |
| Mean Squared Error | 513774655.1 | 539310055 | 537850674.1 |
| Mean Absolute Error | 10864.02602 | 11893.62081 | 11792.14137 |
| R-squared (R^2) | 0.755802044 | 0.743665026 | 0.744358672 |
| Explained Variance Score | 0.756008283 | 0.743687402 | 0.744358817 |
| Max Error | 219424.96 | 244727.712 | 249284.0883 |
| Mean Squared Logarithmic Error | 1.139717029 | 7385.940378 | 7292.788415 |
| Root Mean Squared Error | 22666.59778 | 23223.05008 | 23191.60784 |
| Mean Absolute Percentage Error | 212.1976286 | 221.3735733 | 219.0797703 |

**Figure 3.**
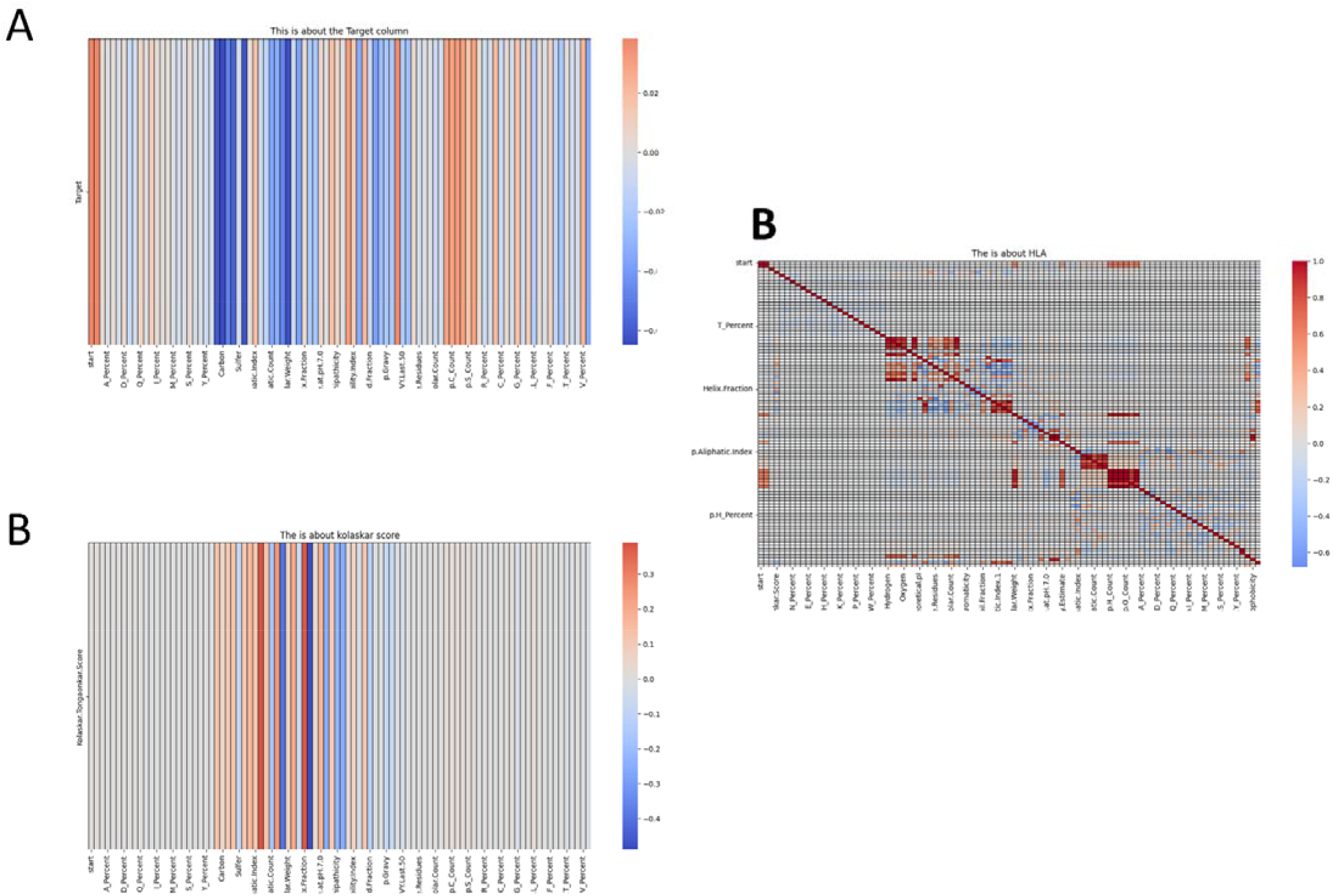
correlation plots of A) Epitope Prediction Model, B) Kolaskar-Tongaonkar(K-T) Score Prediction Model, C) HLA Prediction Model

**Figure 4.**
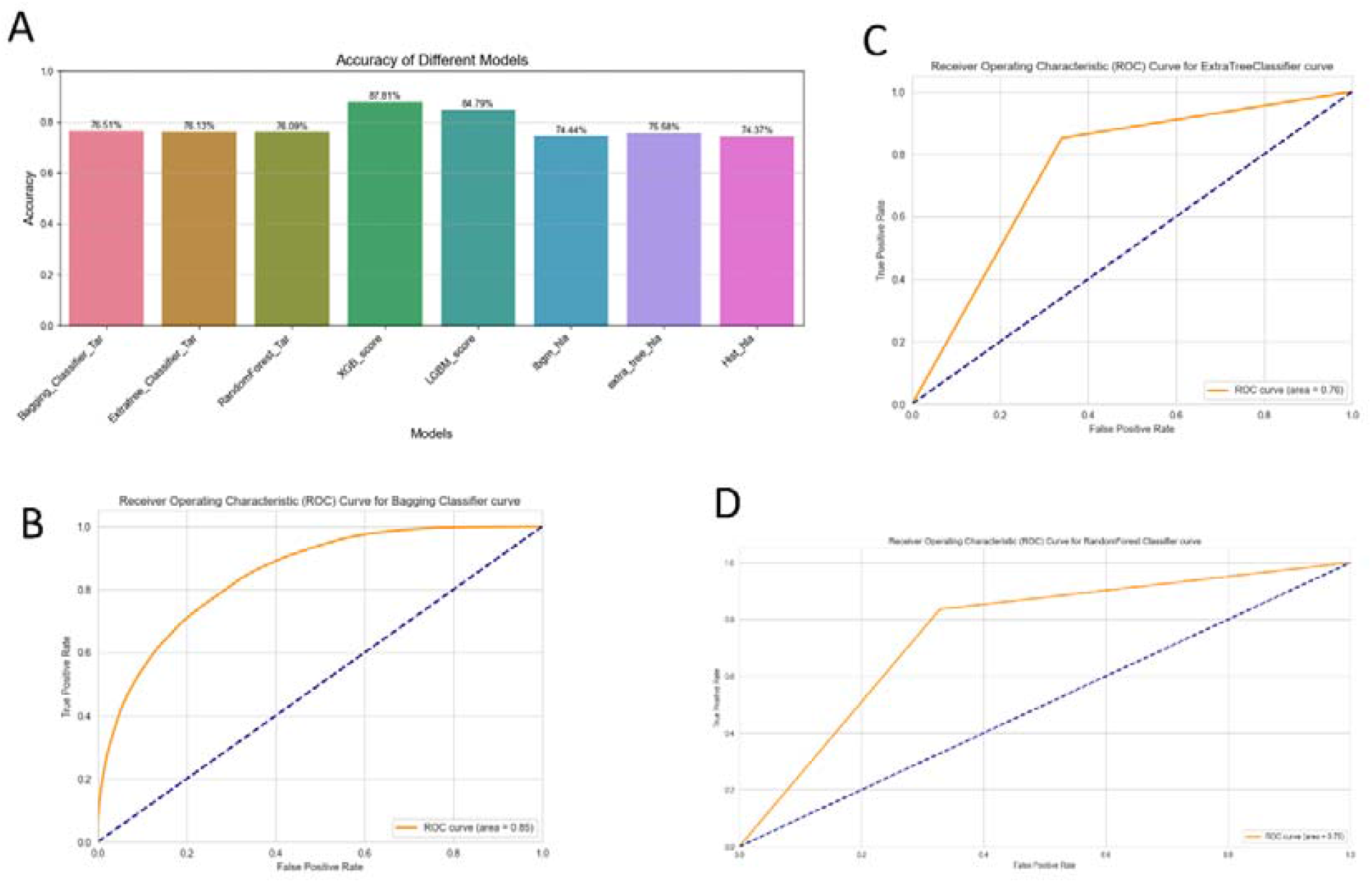
A) Bar charts of accuracy of all the models used in this study, ROC curves of ensemble classifier models used in this study, B) Extra tress classifier, C)Bagging classifier, D)Random forest classifier

#### HLA Prediction Model

In our pursuit to predict HLA epitopes, we initiated with a correlation analysis to discern the most pertinent features within the dataset, extensively outlined in Supplementary Table 2 (Figure 3). Leveraging machine learning, we employed three distinct algorithms: the Extra Trees Regressor, the Hist Gradient Regressor, and the LGBM Regressor[26-28]. To facilitate model comprehension, we engineered features by transforming HLA allele strings into numerical scores via a bespoke Python function, calculate_numerical_score. This innovative approach facilitated the mapping of amino acid sequences from HLA alleles to their respective hydrophobicity values, thereby empowering their inclusion as predictive attributes during model training. Among the trio of algorithms, the LGBM Regressor exhibited superior performance, yielding an impressive R^2 score of 74.44% (as depicted in Figure 4). Renowned for its efficiency and adeptness in handling complex data structures, the LGBM Regressor excelled in capturing intricate relationships within the dataset, affirming its efficacy in accurately predicting HLA epitopes, matric of all models are listed in Table 2.

#### Kolaskar-Tongaonkar(K-T) Score Prediction Model

Our approach to predict K-T scores for epitopes entailed the utilization of backward feature selection, wherein influential features were systematically retained while less informative ones were eliminated. Initially, a correlation analysis was conducted to identify relevant features, providing guidance for the selection process. Features were iteratively removed based on their impact on model performance metrics such as accuracy, precision, recall, and F1-score. Correlated features were meticulously documented in Supplementary Table 2 (Figure 3). For model implementation, we opted for the XGB Regressor and LightGBM Regressor[29-30] due to their effectiveness in capturing complex data patterns. Among these models, the XGB Regressor emerged as the top performer, achieving the highest R^2 score of 87.81% accuracy (as illustrated in Figure 4). Renowned for their capability to handle intricate data relationships, both XGB Regressor and LightGBM Regressor demonstrated their prowess in accurately predicting K-T scores for epitopes, underscoring their suitability for this task, matric of all models are listed in Table 3.

**Table 3.** Results of accuracy and other performance metrics for ensemble regressor model’s (Kolaskar.Tongaonkar.Score prediction).

| Metric | XGB regressor | LightGbm regressor |
| --- | --- | --- |
| Mean Squared Error | 0.000812 | 0.001015 |
| Mean Absolute Error | 0.022459 | 0.025401 |
| R-squared (R^2) | 0.876766 | 0.845912 |
| Explained Variance Score | 0.876791 | 0.845929 |
| Max Error | 0.178647 | 0.183204 |
| Mean Squared Logarithmic Error | 0.000156 | 0.000196 |
| Median Absolute Error | 0.018624 | 0.021538 |
| Mean Poisson Deviance | 0.000635 | 0.000794 |
| Mean Gamma Deviance | 0.000499 | 0.000624 |
| Root Mean Squared Error | 0.028489 | 0.031857 |
| Mean Absolute Percentage Error | 1.760193 | 1.991565 |

#### Comparative Analysis of Models

##### Performance Metrics

Table 1 provides a comprehensive overview of these metrics, including accuracy, AUC (Area Under the Curve), specificity, sensitivity, F-1 score, and precision. Table 1 and Table 3 provide the Mean Squared Error, Mean Absolute Error, R-squared, Explained Variance Score, Max Error, Median Absolute Error, Root Mean Squared Error, and Mean Absolute Percentage Error. These metrics are fundamental in evaluating the models’ ability to classify instances correctly, identify true positives and negatives, minimize false positives and negatives, and maintain a balance between precision and recall[31].

##### Visual Comparison

Figure 4 presents a comparative bar chart illustrating the accuracy of the proposed ensemble model against existing models. This graphical representation offers a quick and intuitive comparison, emphasizing the ensemble model’s dominance. Further insights into the performance of the proposed model are provided through Figure 8, which presents the ROC (Receiver Operating Characteristic) curve[32]. This curve depicts the trade-off between the true positive rate (sensitivity) and the false positive rate (1-specificity) across different threshold values. The AUROC value associated with this curve serves as a quantitative measure of the model’s ability to discriminate between positive and negative instances. In this case, the AUROC value affirms the high discriminatory power of the proposed ensemble model. Overall, the combination of tabular results and visual representations offers a comprehensive evaluation of the proposed ensemble model’s performance, affirming its effectiveness in comparison to standard existing prediction models.

##### Deployment of Trained Models

We’ve taken a crucial step forward in making our epitope prediction models accessible to the public by storing the trained models in .pkl files and deploying them via a Streamlit application hosted on GitHub[33]. This deployment ensures easy access and usability for researchers. Our approach begins by chunking input protein sequences to generate peptide features, distinguishing between epitopes and non-epitopes. Following this, we calculate antigenic scores for the peptides and predict HLAs, culminating in the identification of probable epitopes based on their antigenic scores and other characteristics. Each step in our process is meticulously designed to enhance accuracy and reliability in epitope prediction for cancer immunotherapy. Additionally, we provide visual representations for better understanding and interpretation of the results in Figure 5-A. Through this integrated approach and transparent presentation of results, we aim to accelerate epitope discovery and vaccine design to combat cancer effectively please follow this link for tool (https://vaxoptiml.streamlit.app/).

**Figure 5.**
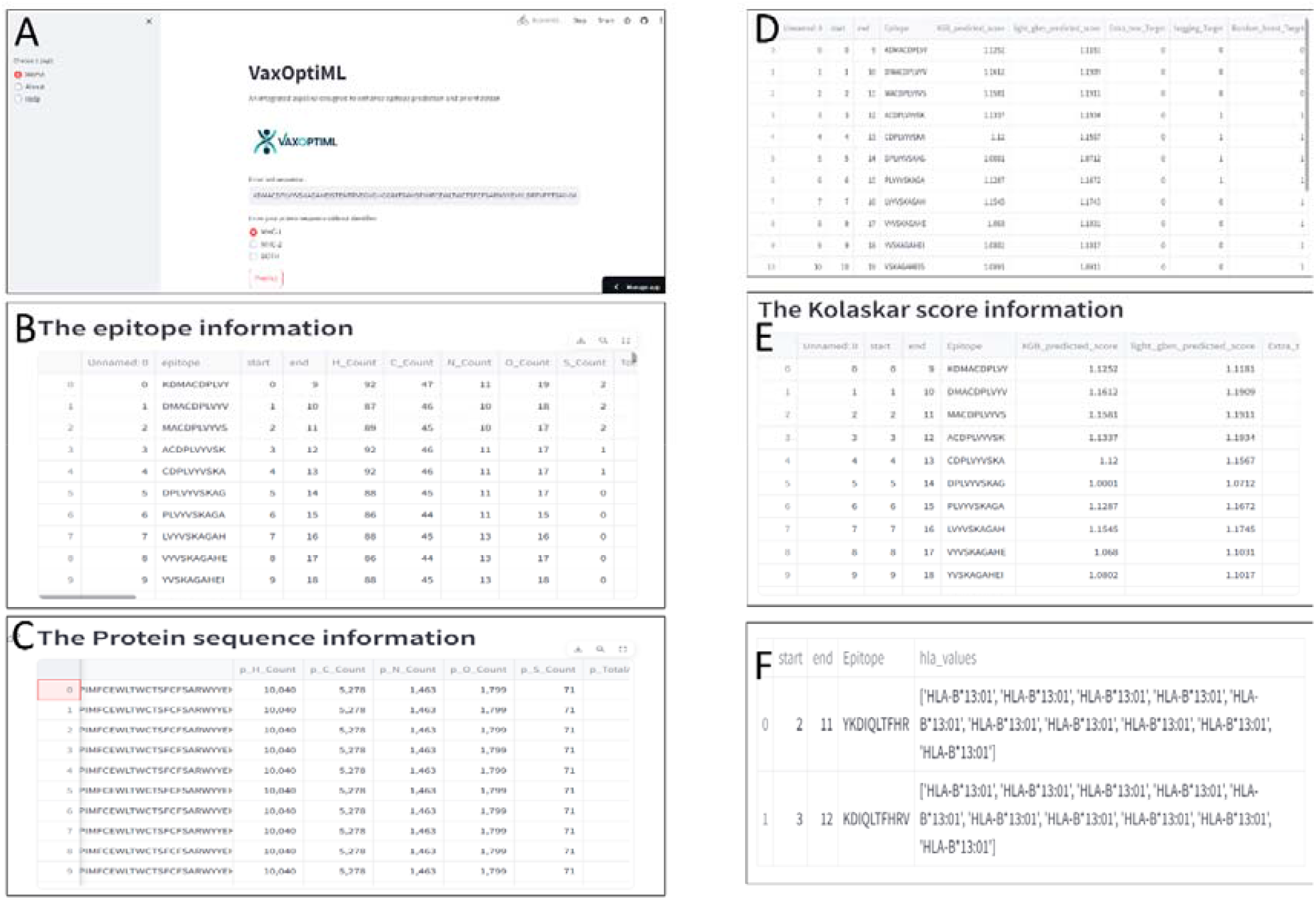
Visual representation of deployed streamlit application, A) Input bar for protein sequence pasting with options, B) Feature extraction for chunked peptide sequence, C) Feature extraction for input protein sequence, D) Probable Epitope through target scores, E) Kolaskar-Tongaonkar(K-T) prediction, F) final output epitope with its binding HLA’s

## Discussion

In the realm of cancer immunotherapy, epitope prediction plays a pivotal role in the development of targeted vaccines, offering a promising avenue for personalized treatment strategies. Cancer, characterized by the uncontrolled growth of abnormal cells, poses significant challenges for traditional therapeutic approaches. However, the advent of immunotherapy has revolutionized cancer treatment by leveraging the body’s immune system to recognize and eliminate tumor cells[34]. Epitopes, short amino acid sequences derived from TAAs, serve as key targets for the immune system’s surveillance and response against cancer. Identifying epitopes that elicit robust immune responses is crucial for designing effective cancer vaccines[35]. Yet, the vast array of potential epitopes, coupled with the complex interplay between tumor cells and the immune system, presents formidable obstacles to epitope prediction. One of the primary challenges in epitope prediction is the heterogeneity of tumor cells. Cancer is not a singular disease but rather a diverse group of diseases characterized by distinct molecular profiles and clinical behaviors[36]. Each tumor type, and even individual tumors within the same type, may harbor unique sets of epitopes, complicating efforts to identify common targets for immunotherapy. Additionally, tumors can undergo genetic mutations and evolve over time, further diversifying the epitope landscape and necessitating ongoing surveillance and adaptation of vaccine strategies. Another challenge in epitope prediction stems from the diversity of HLA alleles in the human population. HLA molecules play a critical role in presenting epitopes to T cells and initiating immune responses. However, different individuals express unique combinations of HLA alleles, resulting in variability in epitope presentation and immune recognition. Predicting epitopes that bind efficiently to diverse HLA alleles is essential for designing vaccines that can elicit robust immune responses across diverse patient populations[37].

Amidst this backdrop, VaxOptiML emerges as a beacon of innovation, offering an integrated pipeline poised to revolutionize epitope prediction and prioritization for cancer vaccines. Anchored by a meticulously curated dataset comprising experimentally validated epitopes and powered by cutting-edge machine learning techniques, VaxOptiML introduces a trifecta of models: epitope prediction, HLA epitope prediction, and Kolaskar-Tongaonkar (K-T) score prediction.

Within the epitope prediction model, the efficacy of ensemble methods takes center stage, with the Bagging Classifier, Extra Trees Classifier, and Random Forest Classifier showcasing notable accuracies of 76.51%, 75.92%, and 74.89%, respectively. These ensemble techniques, celebrated for their ability to mitigate overfitting and enhance predictive robustness, orchestrate a symphony of predictions from diverse models, thereby laying a formidable foundation for the development of targeted cancer vaccines. Transitioning to the HLA epitope prediction model, personalized epitope prediction emerges as a transformative paradigm. Here, the LGBM Regressor rises to prominence, boasting an impressive R^2 score of 74.44%. Personalization is paramount in the realm of cancer immunotherapy, as it navigates the intricate landscape of HLA diversity among individuals. By tailoring epitope predictions to individual patients’ HLA profiles, personalized immunotherapies hold promise for optimizing treatment outcomes and minimizing adverse immune reactions. The accuracy achieved by the LGBM Regressor underscores the significance of leveraging advanced regression algorithms to unravel the complex interplay between epitopes and HLA alleles. Furthermore, the K-T score prediction model unveils its significance in epitope antigenicity prediction, with the XGB Regressor clinching the highest R^2 score of 87.81%. The Kolaskar-Tongaonkar (K-T) score, rooted in the physicochemical properties of amino acid residues, emerges as a critical metric for prioritizing antigenic peptides with high immunogenicity. By accurately forecasting K-T scores for epitopes, researchers can streamline the vaccine design process and expedite the translation of research endeavors into clinical applications. The triumph of the XGB Regressor underscores the efficacy of advanced regression algorithms in epitope antigenicity prediction, heralding a new era in personalized vaccine design.

Over all, VaxOptiML represents a paradigm shift in epitope prediction and epitope-related analyses for cancer immunotherapy. By leveraging curated datasets and advanced machine learning algorithms, VaxOptiML offers a transformative approach to expedite epitope discovery and vaccine design. The robustness and accuracy of its models underscore its potential to significantly impact personalized immunotherapies, paving the way for enhanced treatment outcomes and improved patient care in the fight against cancer. As we continue to push the boundaries of cancer immunology, VaxOptiML stands as a beacon of hope, poised to catalyze innovation and usher in a new era of precision medicine.

## Conclusion

Our study presents a novel approach to epitope prediction and prioritization for cancer immunotherapy, leveraging advanced machine learning techniques and a curated dataset of experimentally validated epitopes. By integrating multiple models targeting epitope prediction, personalized HLA pairing, and immunogenicity prioritization, we provide a comprehensive framework for accelerating the identification of promising vaccine candidates. Through rigorous evaluation and comparison with existing prediction models, we demonstrate the superior performance of our ensemble model, highlighting its potential to revolutionize epitope discovery processes in cancer immunotherapy. Moving forward, our approach holds promise for driving advancements in personalized medicine and contributing to the development of more effective immunotherapeutic strategies for cancer treatment.

## Supporting information

https://docs.google.com/document/d/1pwElZDN1i4FfztL09ECjvcT9Lo-DVQst/edit#heading=h.gjdgxs

## Acknowledgments

The authors express deep gratitude to the management of REVA University, Vellore Institute of Technology and Institute of bioinformatics (IOB) for providing all the support, necessary facilities, assistance, and constant encouragement to carry out this work.

## Data Availability Statement

All the codes and data used for this study are available in https://github.com/karthick1087/VaxOptiML

