## Supplementary material for "VaxOptiML: Leveraging Machine Learning for Accurate Prediction of MHC-I & II Epitopes for Optimized Cancer Immunotherapy": https://docs.google.com/document/d/1pwElZDN1i4FfztL09ECjvcT9Lo-DVQst/edit#heading=h.gjdgxs

*Corresponding

Karthick Vasudevan, Institute of Bioinformatics, International Technology Park, Bangalore, 560066, India.

Supplementary Table 1. Extracted features of epitopes and protein sequence

| Sl.no | features of epitopes | features of epitopes |
| --- | --- | --- |
| 1 | Kolaskar.Tongaonkar.Score | p.Molecular.Weight |
| 2 | A_Percent | p.Instability.Index |
| 3 | R_Percent | p.Aromaticity |
| 4 | N_Percent | p.Helix.Fraction |
| 5 | D_Percent | p.Strand.Fraction |
| 6 | C_Percent | p.Coil.Fraction |
| 7 | E_Percent | p.Charge.at.pH.7.0 |
| 8 | Q_Percent | p.Gravy |
| 9 | G_Percent | p.Aliphatic.Index.1 |
| 10 | H_Percent | p.Amphipathicity.Estimate |
| 11 | I_Percent | p.GRAVY.Last.50 |
| 12 | L_Percent | p.Molar.Extinction.Coefficient |
| 13 | K_Percent | p.Theoretical.pI |
| 14 | M_Percent | p.Aliphatic.Index |
| 15 | F_Percent | p.Positive.Residues |
| 16 | P_Percent | p.Negative.Residues |
| 17 | S_Percent | p.Aromatic.Count |
| 18 | T_Percent | p.Polar.Count |
| 19 | W_Percent | p.Nonpolar.Count |
| 20 | Y_Percent | p.H_Count |
| 21 | V_Percent | p.C_Count |
| 22 | Hydrogen | p.N_Count |
| 23 | Carbon | p.O_Count |
| 24 | Nitrogen | p.S_Count |
| 25 | Oxygen | p.TotalAtoms_Count |
| 26 | Sulfer | p.A_Percent |
| 27 | TotalAtoms | p.R_Percent |
| 28 | Theoretical.pI | p.N_Percent |
| 29 | Aliphatic.Index | p.D_Percent |
| 30 | Positive.Residues | p.C_Percent |
| 31 | Negative.Residues | p.E_Percent |
| 32 | Aromatic.Count | p.Q_Percent |
| 33 | Polar.Count | p.G_Percent |
| 34 | Nonpolar.Count | p.H_Percent |
| 35 | Molecular.Weight | p.I_Percent |
| 36 | Instability.Index | p.L_Percent |
| 37 | Aromaticity | p.K_Percent |
| 38 | Helix.Fraction | p.M_Percent |
| 39 | Strand.Fraction | p.F_Percent |
| 40 | Coil.Fraction | p.P_Percent |
| 41 | Charge.at.pH.7.0 | p.S_Percent |
| 42 | Gravy | p.T_Percent |
| 43 | Aliphatic.Index.1 | p.W_Percent |
| 44 | Amphipathicity | p.Y_Percent |
| 45 | GRAVY.Last.50 | p.V_Percent |
| 46 | Molar.Extinction.Coefficient |  |

Supplementary Table 2. Correlating features of all the 3 ml models

| Model for epitope prediction based on target | Model for HLA prediction for predicted epitopes | Model for Kolaskar.Tongaonkar.Score prediction for predicted epitopes |
| --- | --- | --- |
| start | start | Target |
| end | end | C_Percent |
| R_Percent | A_Percent | Q_Percent |
| D_Percent | R_Percent | G_Percent |
| Q_Percent | N_Percent | K_Percent |
| I_Percent | D_Percent | P_Percent |
| L_Percent | C_Percent | S_Percent |
| S_Percent | E_Percent | T_Percent |
| Aliphatic.Index | Q_Percent | W_Percent |
| Helix.Fraction | G_Percent | H_Count |
| Aliphatic.Index.1 | H_Percent | C_Count |
| p.Molecular.Weight | I_Percent | N_Count |
| p.Instability.Index | L_Percent | O_Count |
| p.Amphipathicity.Estimate | K_Percent | TotalAtoms_Count |
| p.H_Count | M_Percent | Theoretical.pI |
| p.N_Count | F_Percent | Positive.Residues |
| p.S_Count | P_Percent | Negative.Residues |
| p.TotalAtoms_Count | S_Percent | Aromatic.Count |
| p.D_Percent | T_Percent | Polar.Count |
| p.E_Percent | W_Percent | Nonpolar.Count |
| p.I_Percent | Y_Percent | Molecular.Weight |
| p.F_Percent | V_Percent | Instability.Index |
| p.V_Percent | Hydrogen | Strand.Fraction |
| H_Percent | Carbon | Charge.at.pH.7.0 |
| K_Percent | Nitrogen | p_Aromaticity |
| Theoretical.pI | Sulfer | p_Strand.Fraction |
| Charge.at.pH.7.0 | TotalAtoms | p_Coil.Fraction |
| Amphipathicity | Theoretical.pI | p_Gravy |
| p.Helix.Fraction | Aliphatic.Index | p_Amphipathicity |
| p.Aliphatic.Index | Positive.Residues | p_GRAVY.Last.50 |
| p.C_Count | Negative.Residues | p_Aliphatic.Index |
| p.O_Count | Aromatic.Count | p_Polar.Count |
| p.A_Percent | Polar.Count | p_N_Percent |
| p.G_Percent | Nonpolar.Count | p_C_Percent |
| p.K_Percent | Molecular.Weight | p_K_Percent |
| p.T_Percent | Instability.Index | p_F_Percent |
|  | Aromaticity | p_P_Percent |
|  | Helix.Fraction | p_S_Percent |
|  | Strand.Fraction | p_T_Percent |
|  | Coil.Fraction | p_W_Percent |
|  | Charge.at.pH.7.0 | p_Y_Percent |
|  | Amphipathicity | p_V_Percent |
|  | GRAVY.Last.50 | type |
|  | p.Instability.Index | hydrophobicity |
|  | p.Helix.Fraction |  |
|  | p.Strand.Fraction |  |
|  | p.Coil.Fraction |  |
|  | p.Charge.at.pH.7.0 |  |
|  | p.Amphipathicity.Estimate |  |
|  | p.Aliphatic.Index |  |
|  | p.Aromatic.Count |  |
|  | p.Nonpolar.Count |  |
|  | p.H_Count |  |
|  | p.C_Count |  |
|  | p.O_Count |  |
|  | p.TotalAtoms_Count |  |
|  | p.R_Percent |  |
|  | p.N_Percent |  |
|  | p.D_Percent |  |
|  | p.E_Percent |  |
|  | p.L_Percent |  |
|  | p.T_Percent |  |
